## Supplemental figures 1-10 for "Long-read single-cell sequencing reveals expressions of hypermutation clusters of isoforms in human liver cancer cells"

### Slide 1
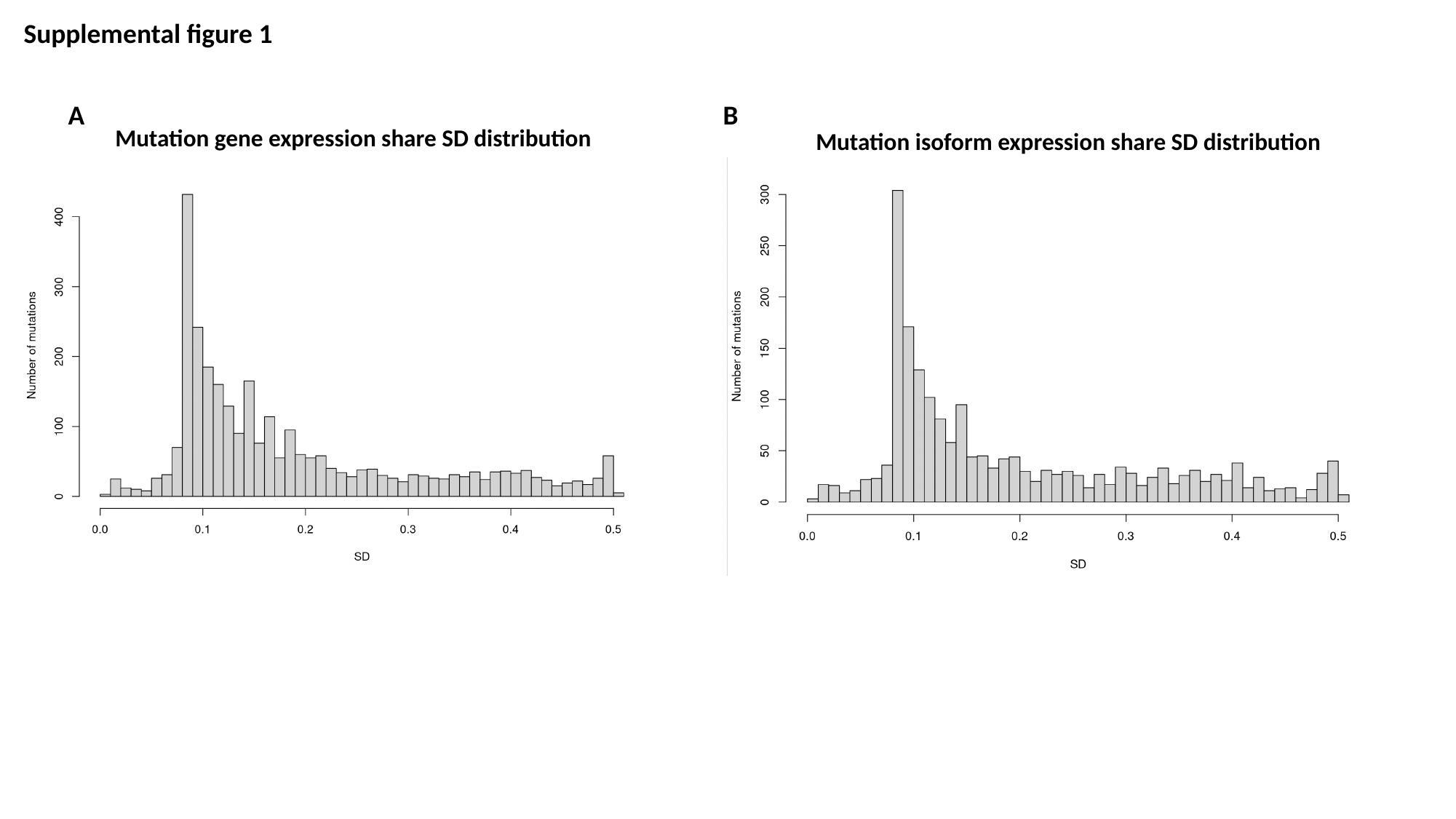

Supplemental figure 1
A						B
Mutation gene expression share SD distribution
Mutation isoform expression share SD distribution

### Slide 2
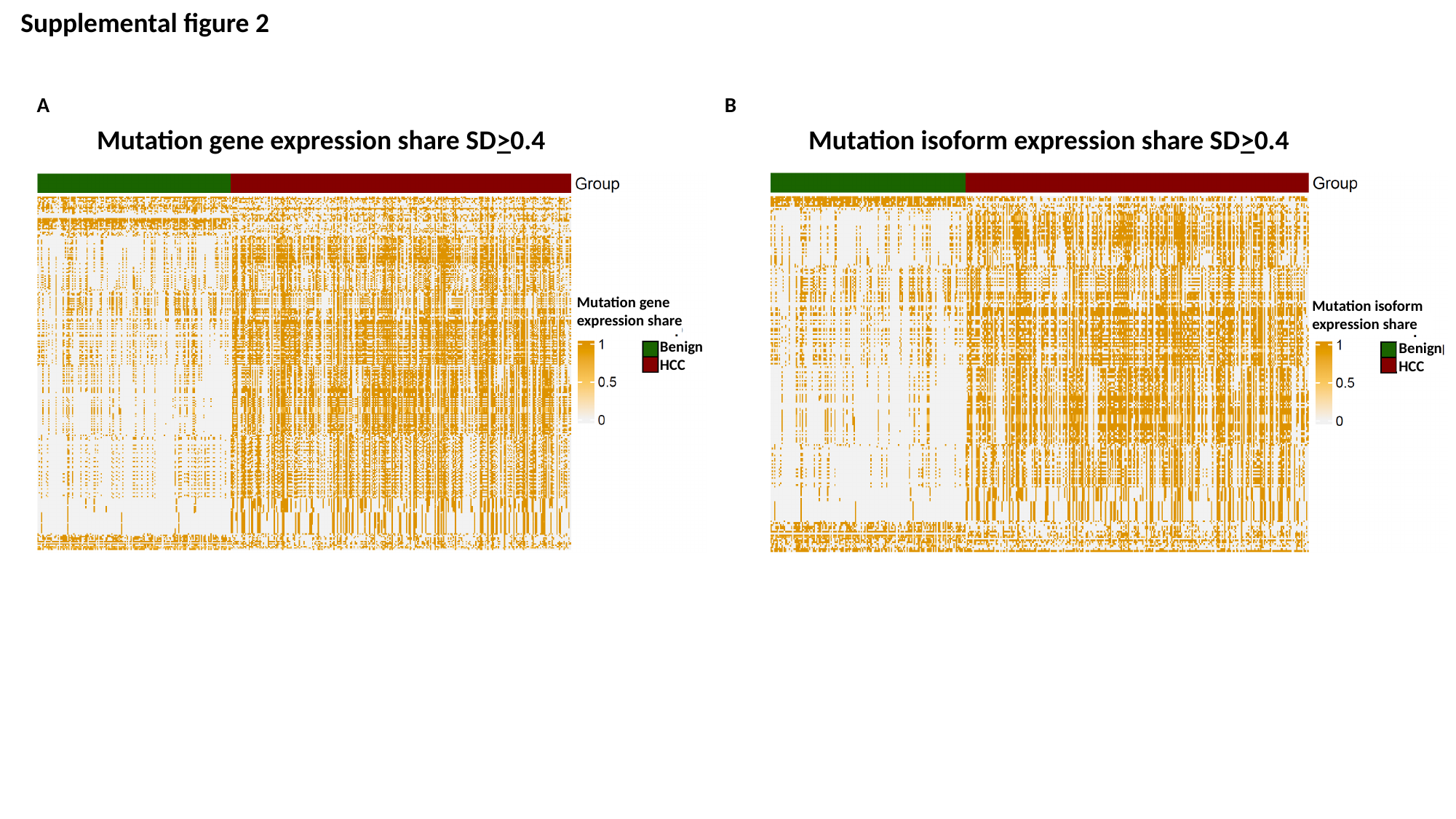

Supplemental figure 2
A						 B
Mutation gene expression share SD>0.4
Mutation isoform expression share SD>0.4
Mutation gene
expression share
Mutation isoform
expression share
Benign
HCC
Benign
HCC

### Slide 3
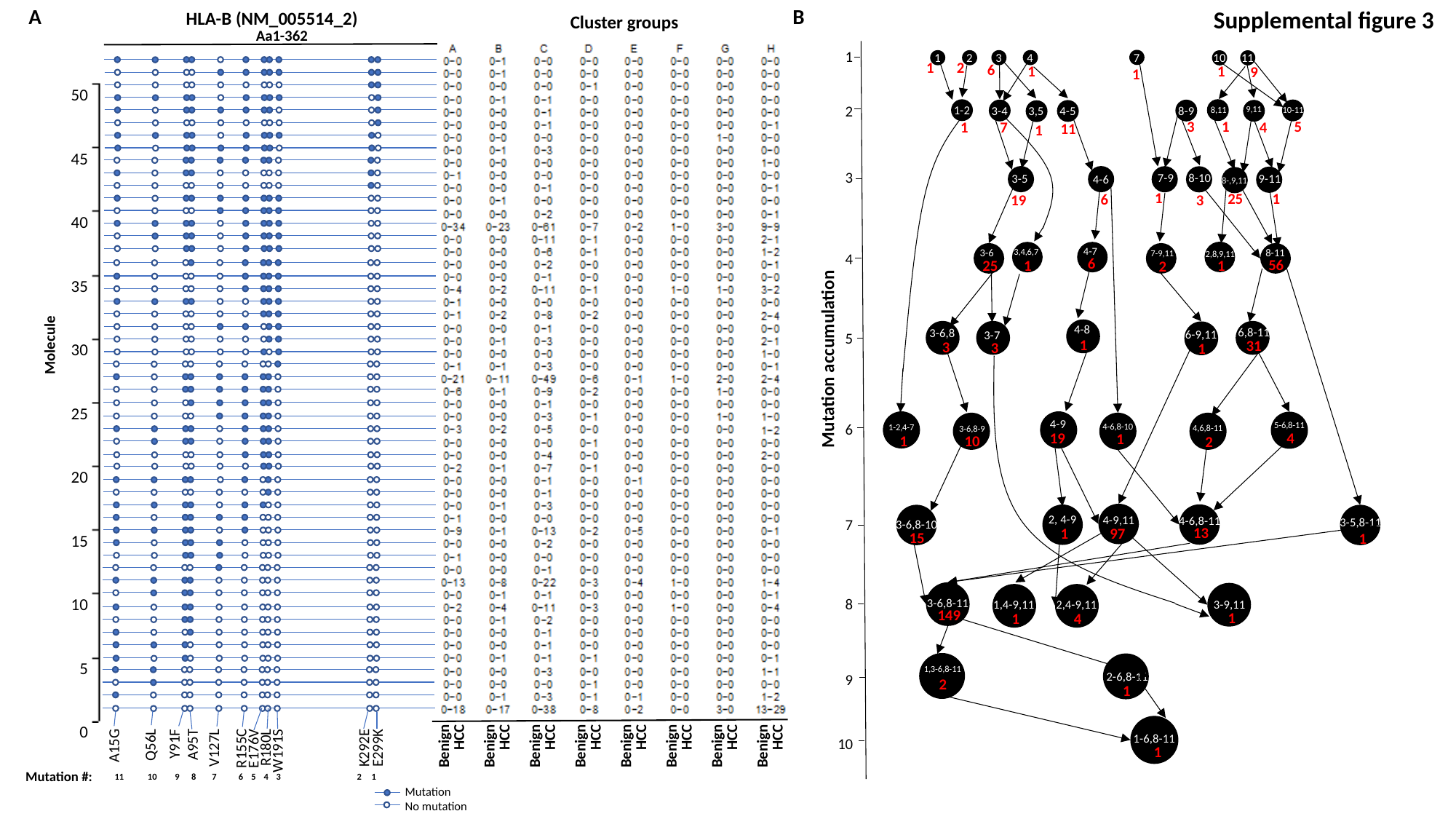

A							B
Supplemental figure 3
HLA-B (NM_005514_2)
Cluster groups
Aa1-362
50
45
40
35
30
25
20
15
10
5
0
1
2
3
4
5
6
7
8
9
10
7
4
3
1
2
10
11
2
1
6
1
1
9
1
1-2
3-4
8-9
9,11
4-5
3,5
10-11
8,11
1
5
3
7
4
1
11
1
7-9
8-10
9-11
3-5
4-6
8-,9,11
1
25
1
6
3
19
4-7
3-6
8-11
3,4,6,7
2,8,9,11
7-9,11
6
56
25
1
1
2
4-8
6,8-11
3-6,8
3-7
6-9,11
1
Molecule
31
3
3
1
Mutation accumulation
4-9
1-2,4-7
5-6,8-11
3-6,8-9
4-6,8-10
4,6,8-11
19
4
1
1
10
2
4-9,11
4-6,8-11
2, 4-9
3-6,8-10
3-5,8-11
13
97
1
15
1
Benign
HCC
Benign
HCC
Benign
HCC
Benign
HCC
Benign
HCC
Benign
HCC
Benign
HCC
Benign
HCC
3-9,11
1,4-9,11
2,4-9,11
A15G
Q56L
Y91F
A95T
V127L
R155C
E176V
R180L
W191S
K292E
E299K
3-6,8-11
149
1
1
4
1,3-6,8-11
2-6,8-11
2
1
1-6,8-11
1
Mutation #:
11 10 9 8 7 6 5 4 3 2 1
Mutation
No mutation

### Slide 4
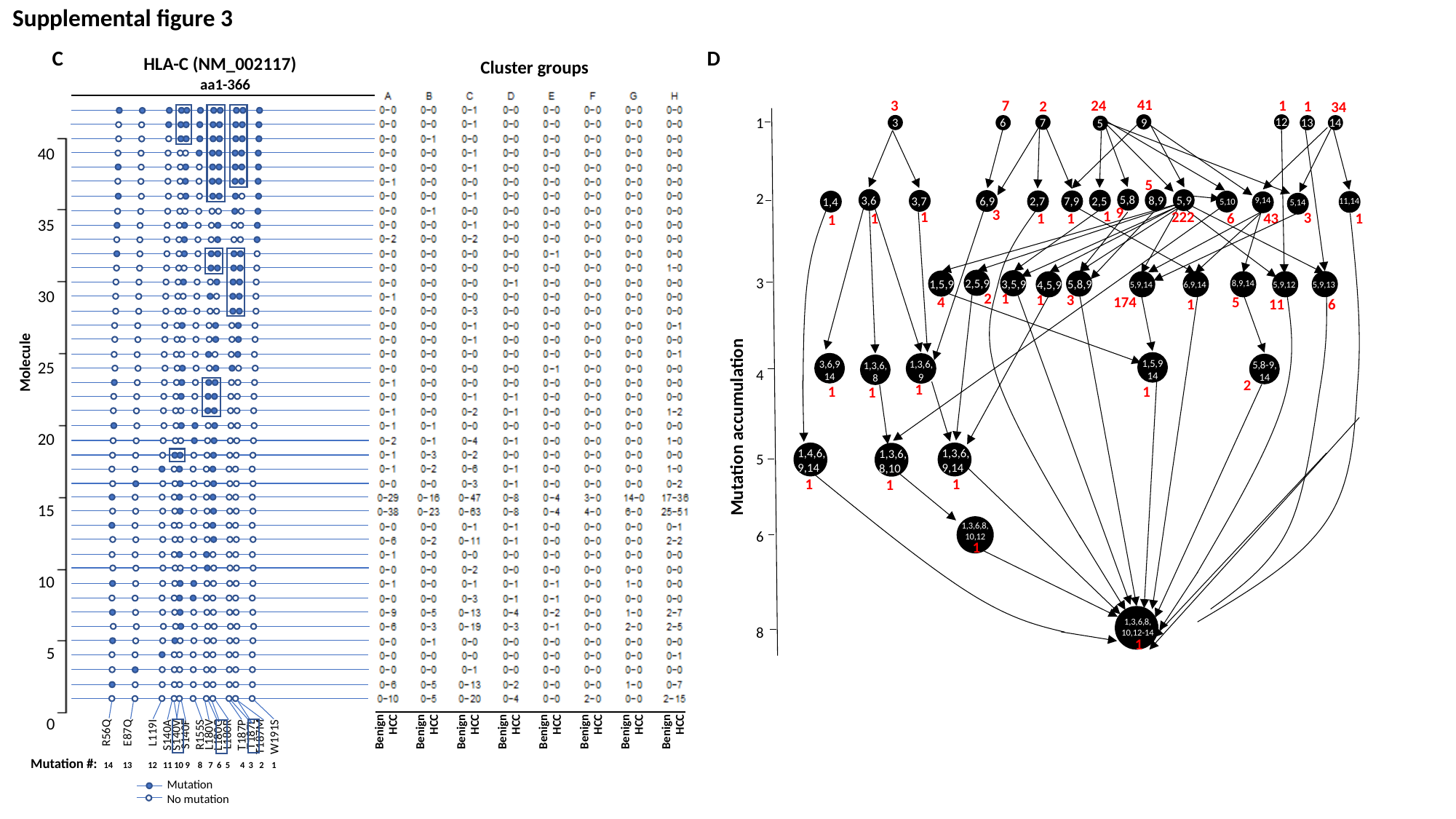

Supplemental figure 3
C						D
HLA-C (NM_002117)
Cluster groups
aa1-366
40
35
30
25
20
15
10
5
0
41
7
3
24
1
2
1
34
1
2
3
4
5
6
8
12
7
3
6
9
13
14
5
5
5,8
3,6
8,9
5,9
2,5
3,7
6,9
2,7
7,9
1,4
9,14
11,14
5,10
5,14
9
3
1
1
222
3
1
1
1
6
43
1
1
2,5,9
3,5,9
1,5,9
5,8,9
6,9,14
5,9,13
5,9,14
8,9,14
5,9,12
4,5,9
1
2
3
1
4
174
5
6
11
1
Molecule
1,5,9
14
3,6,9
14
1,3,6,
9
5,8-9,
14
1,3,6,
8
2
1
1
1
1
Mutation accumulation
1,4,6,
9,14
1,3,6,
9,14
1,3,6,
8,10
1
1
1
1,3,6,8,
10,12
1
Benign
HCC
Benign
HCC
Benign
HCC
Benign
HCC
Benign
HCC
Benign
HCC
Benign
HCC
Benign
HCC
1,3,6,8,
10,12-14
1
R56Q
E87Q
L119I
S140A
S140V
S140F
R155S
L180V
L180G
L180R
T187P
T187L
T187M
W191S
Mutation #:
14 13 12 11 10 9 8 7 6 5 4 3 2 1
Mutation
No mutation

### Slide 5
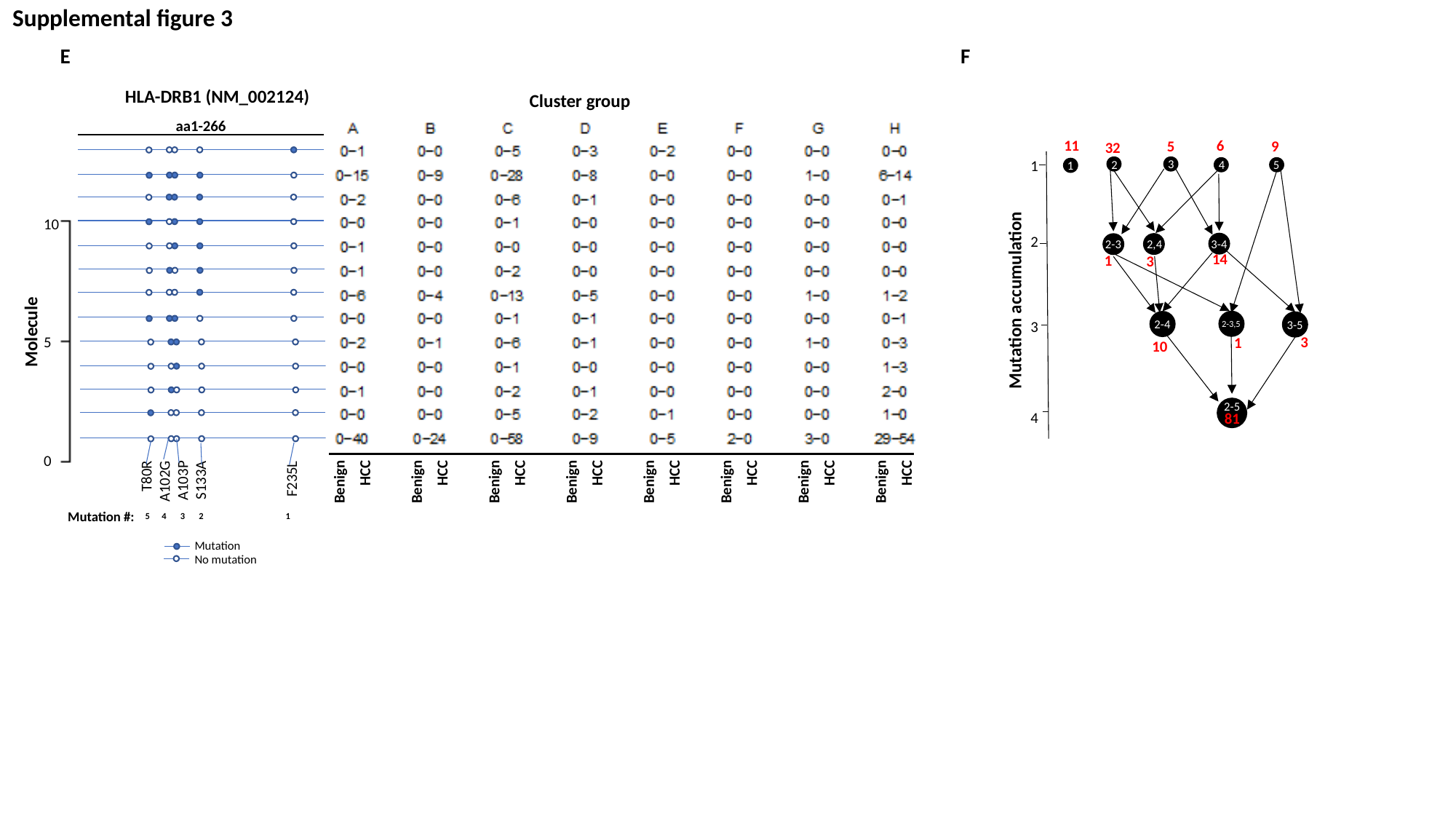

Supplemental figure 3
E						 F
HLA-DRB1 (NM_002124)
Cluster group
aa1-266
6
11
9
5
32
3
1
2
3
4
2
5
4
1
Benign
HCC
Benign
HCC
Benign
HCC
Benign
HCC
Benign
HCC
Benign
HCC
Benign
HCC
Benign
HCC
10
5
0
3-4
2,4
2-3
14
1
3
Mutation accumulation
2-3,5
2-4
3-5
Molecule
3
1
10
T80R
A102G
A103P
S133A
F235L
2-5
81
Mutation #:
5 4 3 2 1
Mutation
No mutation

### Slide 6
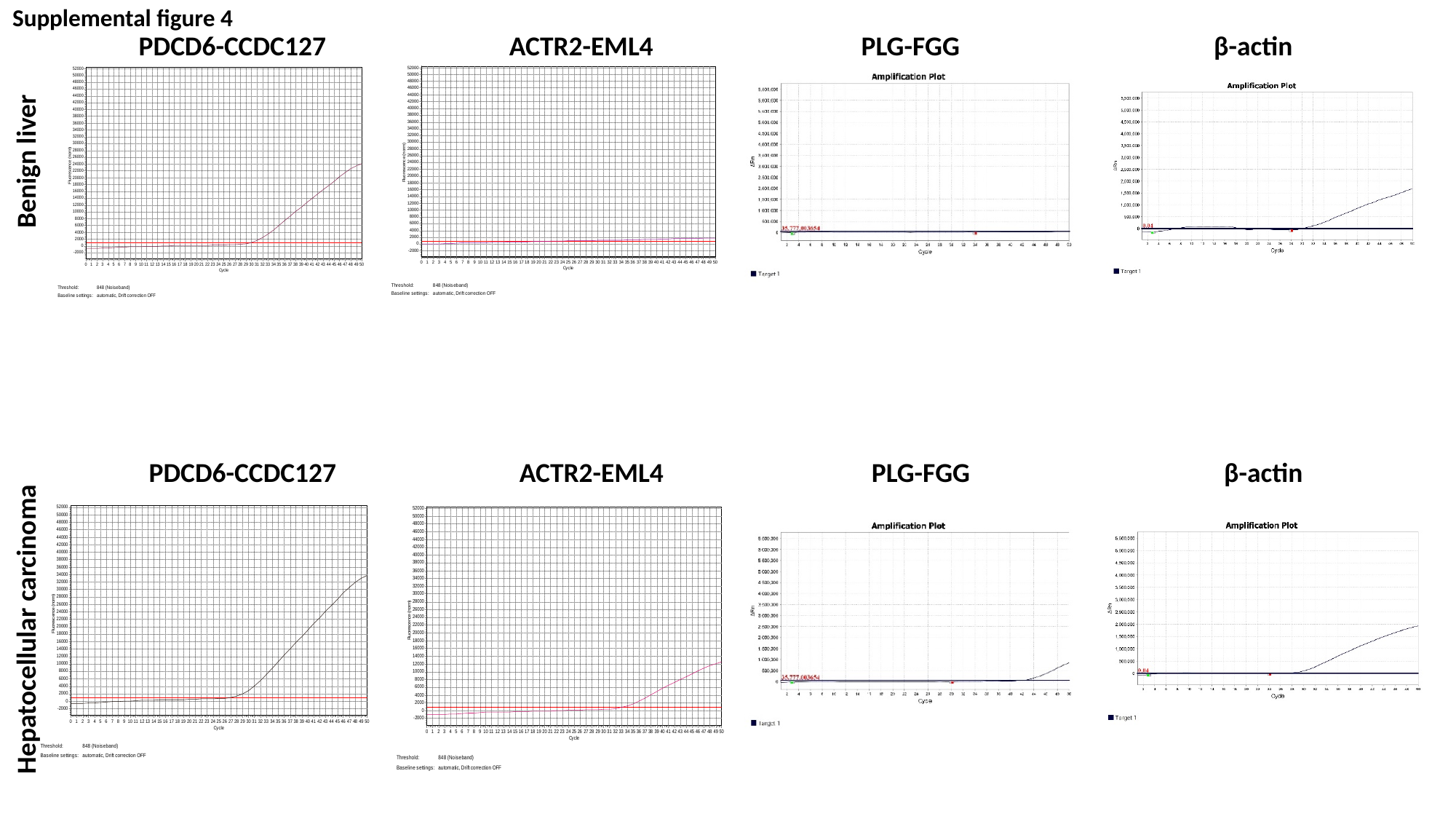

Supplemental figure 4
PDCD6-CCDC127		 ACTR2-EML4		 PLG-FGG		 β-actin
Hepatocellular carcinoma			Benign liver
PDCD6-CCDC127		 ACTR2-EML4		 PLG-FGG		 β-actin

### Slide 7
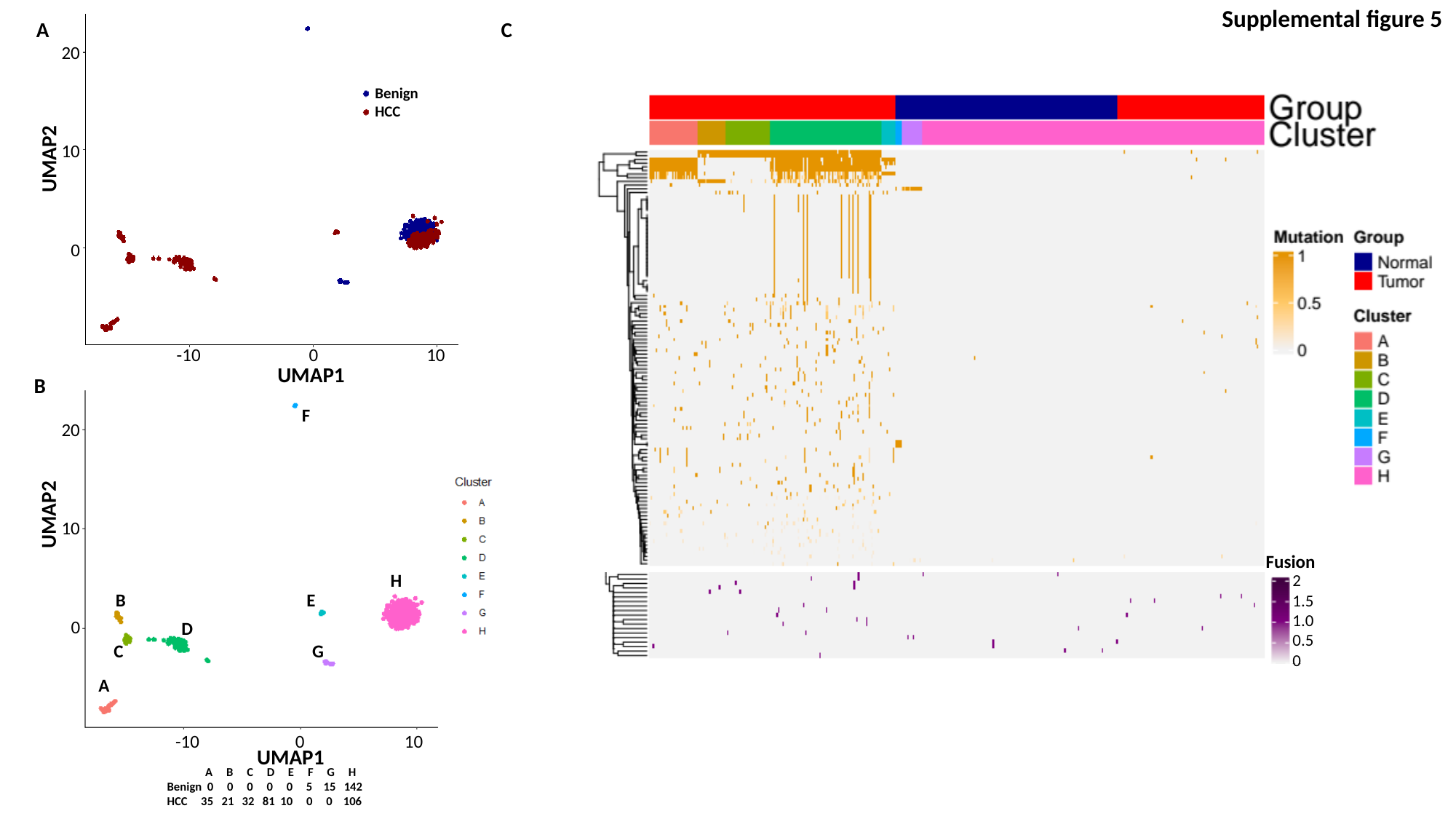

Supplemental figure 5
A				 C
20
10
0
Benign
HCC
UMAP2
-10 0 10
UMAP1
B
G
F
20
10
0
UMAP2
Fusion
H
H
2
1.5
1.0
0.5
0
E
B
E
B
D
D
C
C
F
G
A
A
-10 0 10
UMAP1
 A B C D E F G H
Benign 0 0 0 0 0 5 15 142
HCC 35 21 32 81 10 0 0 106

### Slide 8
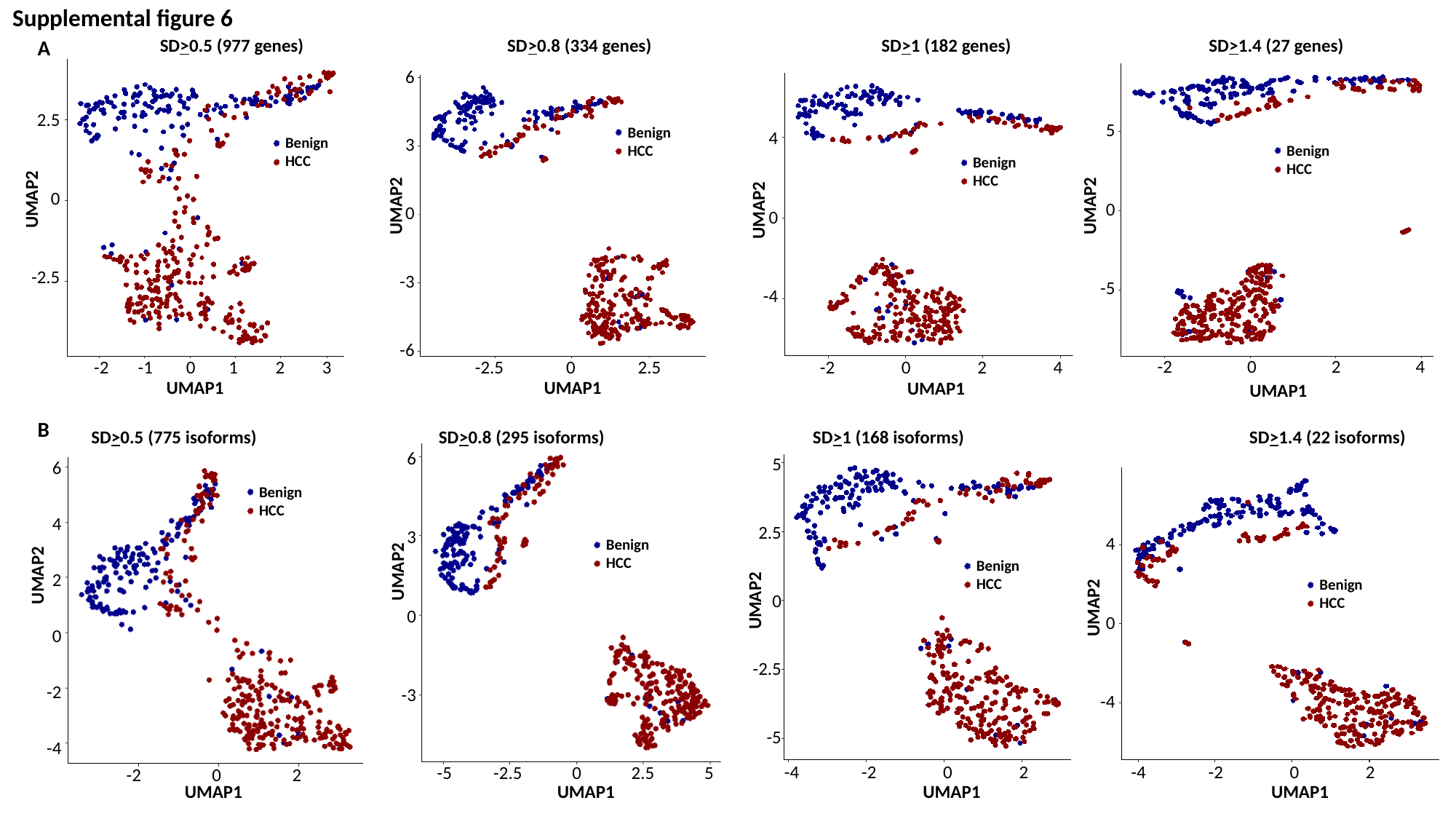

Supplemental figure 6
A
B
SD>0.5 (977 genes)	 SD>0.8 (334 genes)		 SD>1 (182 genes)		 SD>1.4 (27 genes)
6
3
0
-3
-6
2.5
0
-2.5
5
0
-5
Benign
HCC
4
0
-4
Benign
HCC
Benign
HCC
Benign
HCC
UMAP2
UMAP2
UMAP2
UMAP2
-2 0 2 4
-2 -1 0 1 2 3
-2.5 0 2.5
-2 0 2 4
UMAP1
UMAP1
UMAP1
UMAP1
SD>0.5 (775 isoforms)	 SD>0.8 (295 isoforms)		 SD>1 (168 isoforms)		 SD>1.4 (22 isoforms)
6
3
0
-3
5
2.5
0
-2.5
-5
6
4
2
0
-2
-4
Benign
HCC
4
0
-4
Benign
HCC
Benign
HCC
UMAP2
UMAP2
Benign
HCC
UMAP2
UMAP2
-4 -2 0 2
-4 -2 0 2
-5 -2.5 0 2.5 5
-2 0 2
UMAP1		 UMAP1			 UMAP1		 UMAP1

### Slide 9
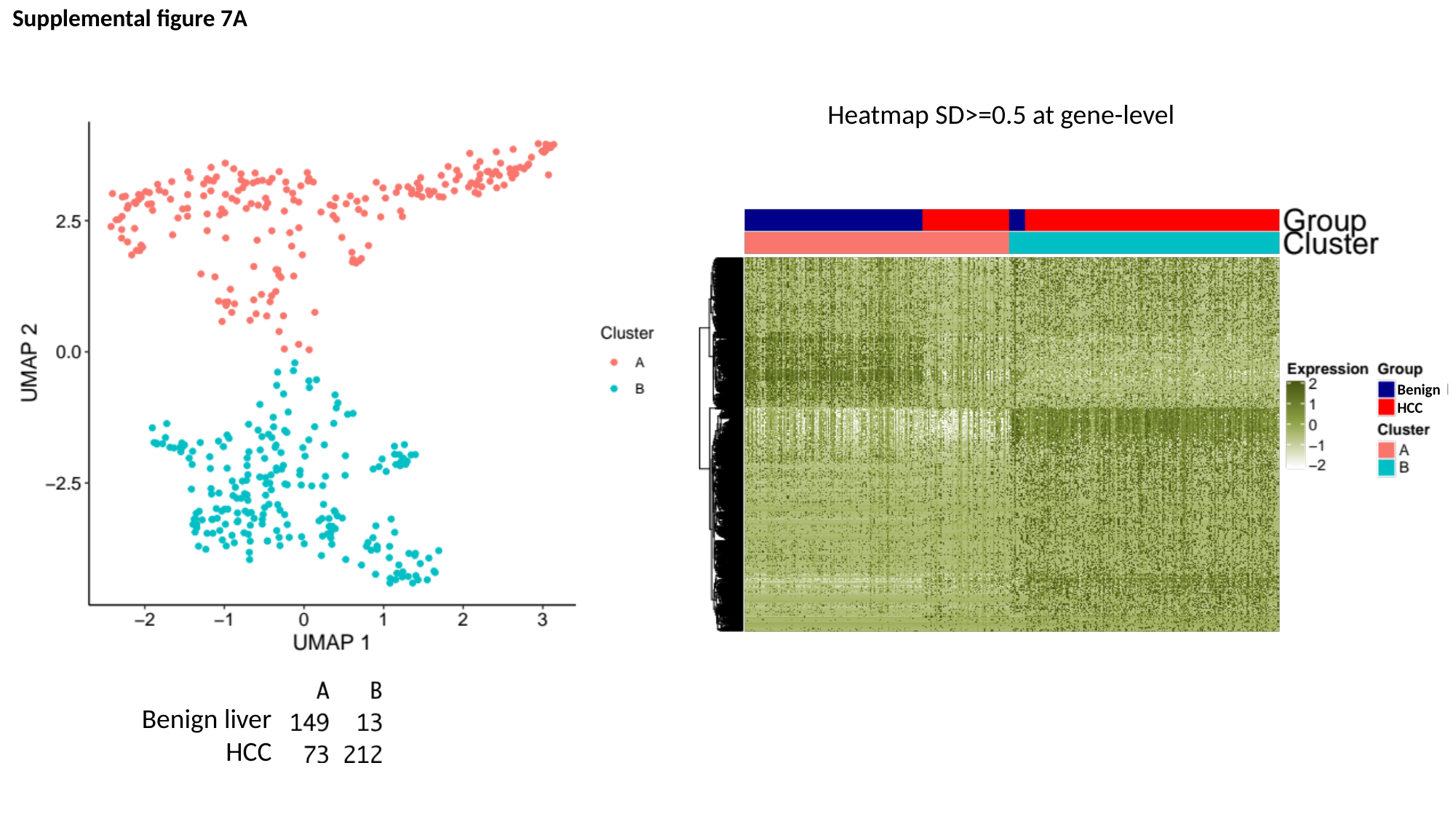

Supplemental figure 7A
Heatmap SD>=0.5 at gene-level
Benign
HCC
Benign liver
HCC

### Slide 10
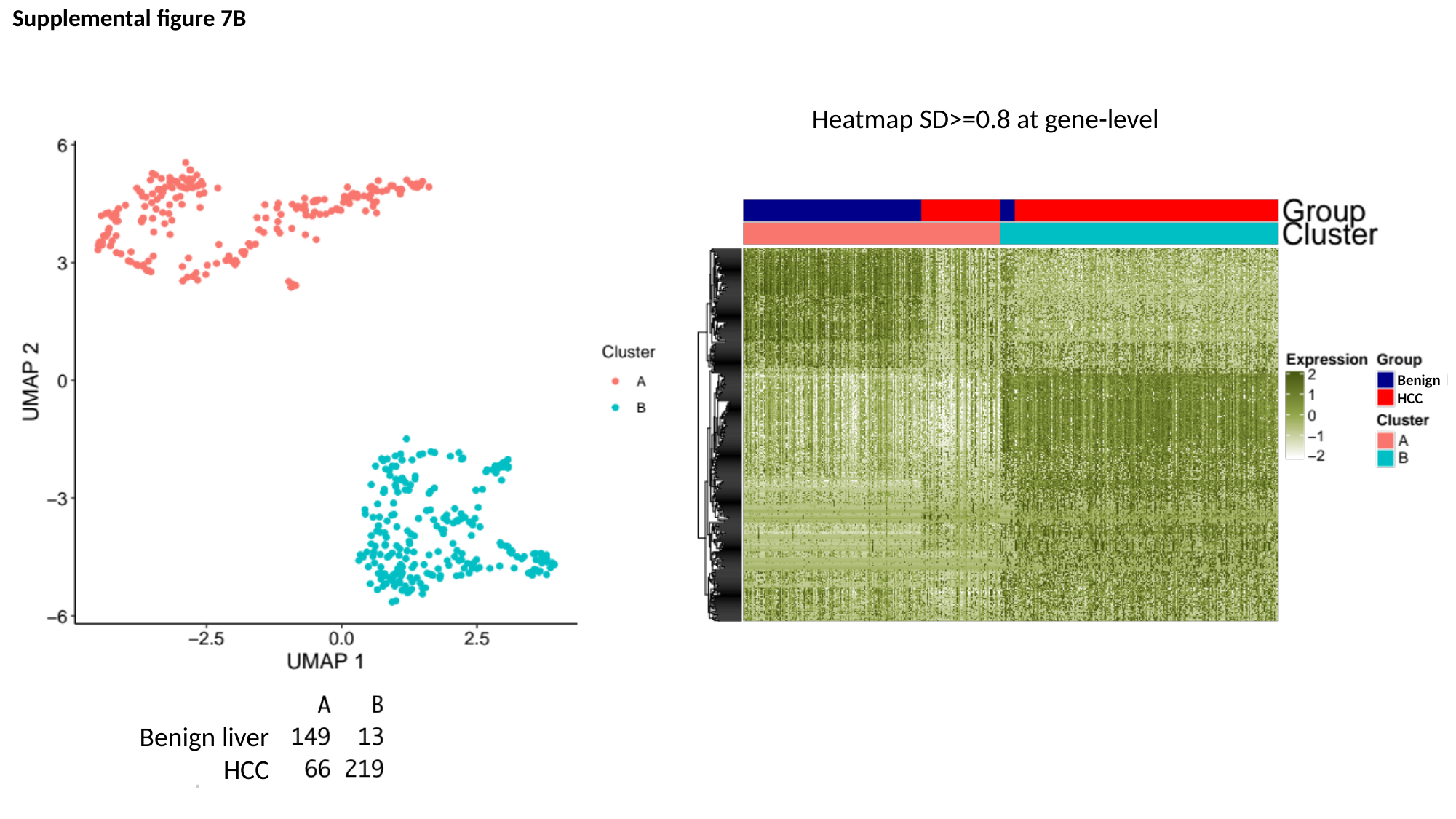

Supplemental figure 7B
Heatmap SD>=0.8 at gene-level
Benign
HCC
Benign liver
HCC

### Slide 11
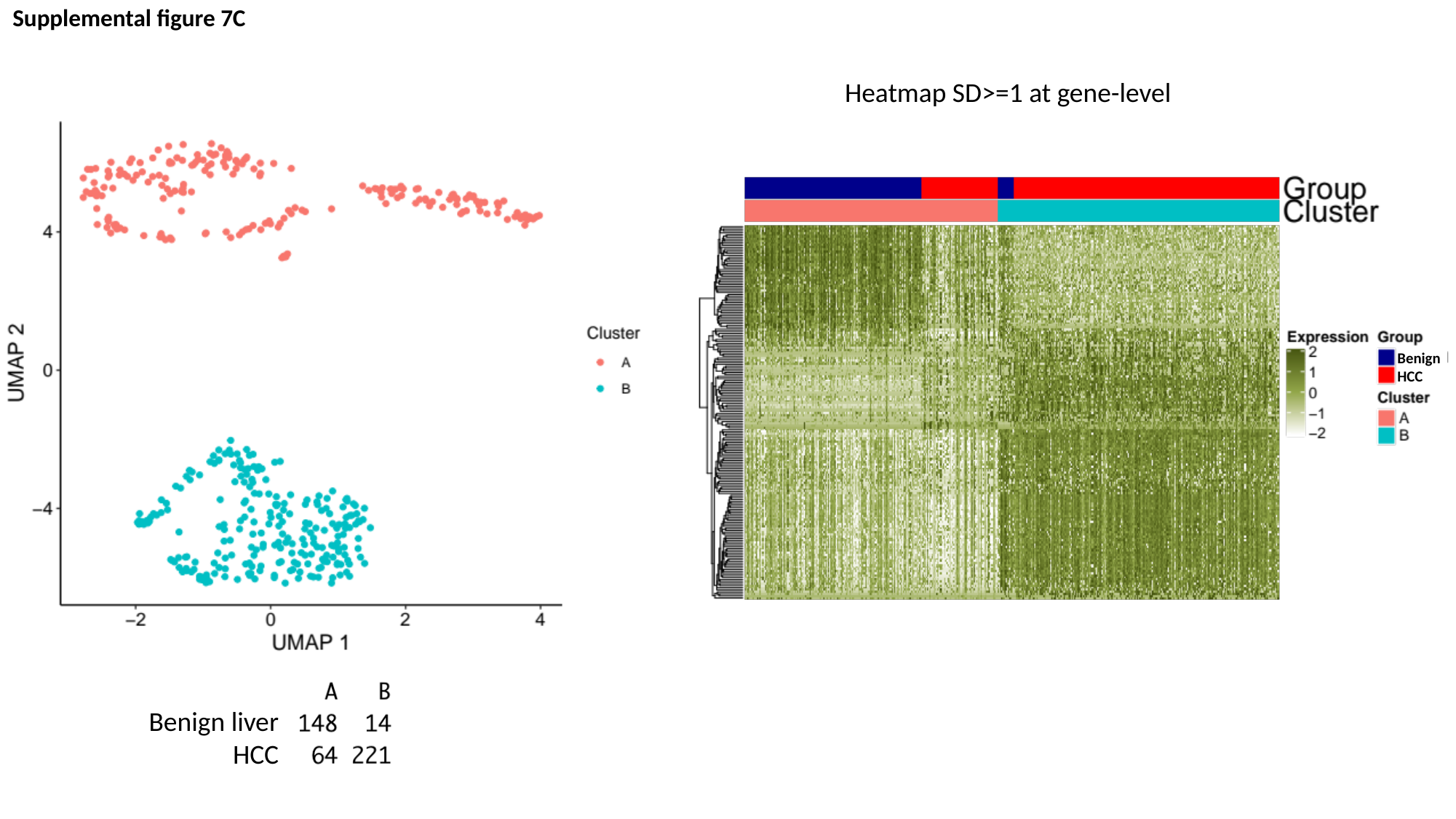

Supplemental figure 7C
Heatmap SD>=1 at gene-level
Benign
HCC
Benign liver
HCC

### Slide 12
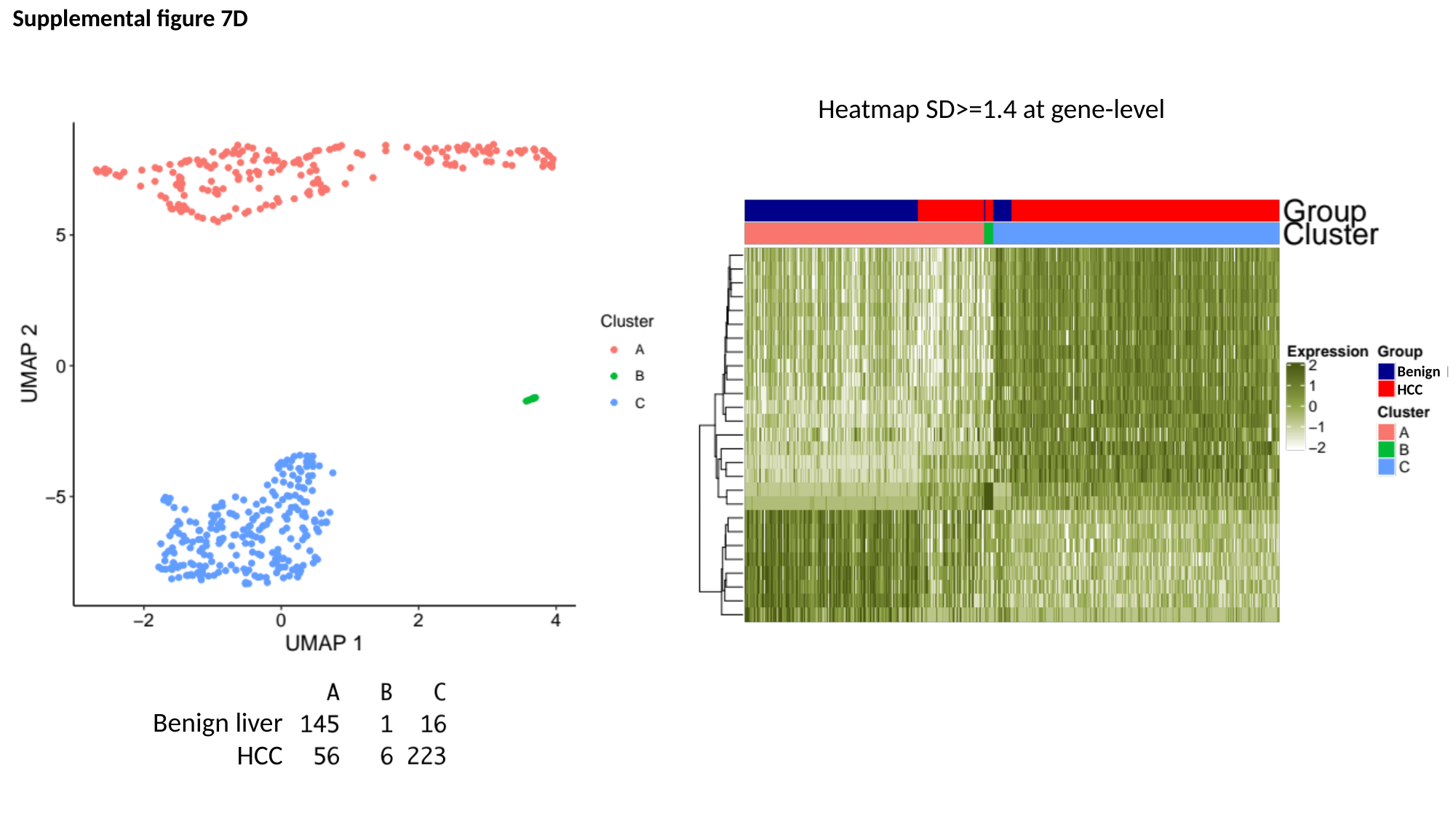

Supplemental figure 7D
Heatmap SD>=1.4 at gene-level
Benign
HCC
Benign liver
HCC

### Slide 13
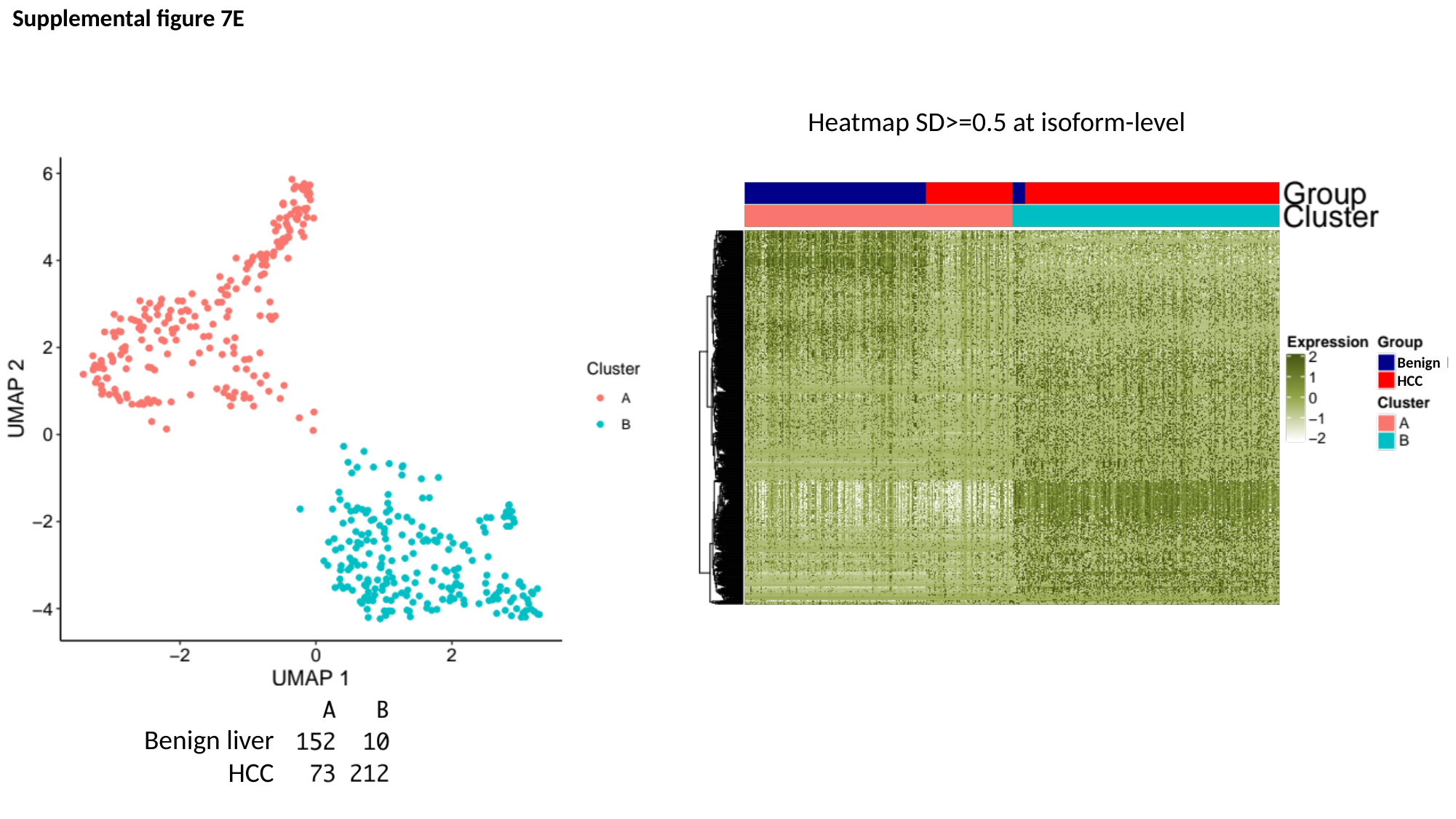

Supplemental figure 7E
Heatmap SD>=0.5 at isoform-level
Benign
HCC
Benign liver
HCC

### Slide 14
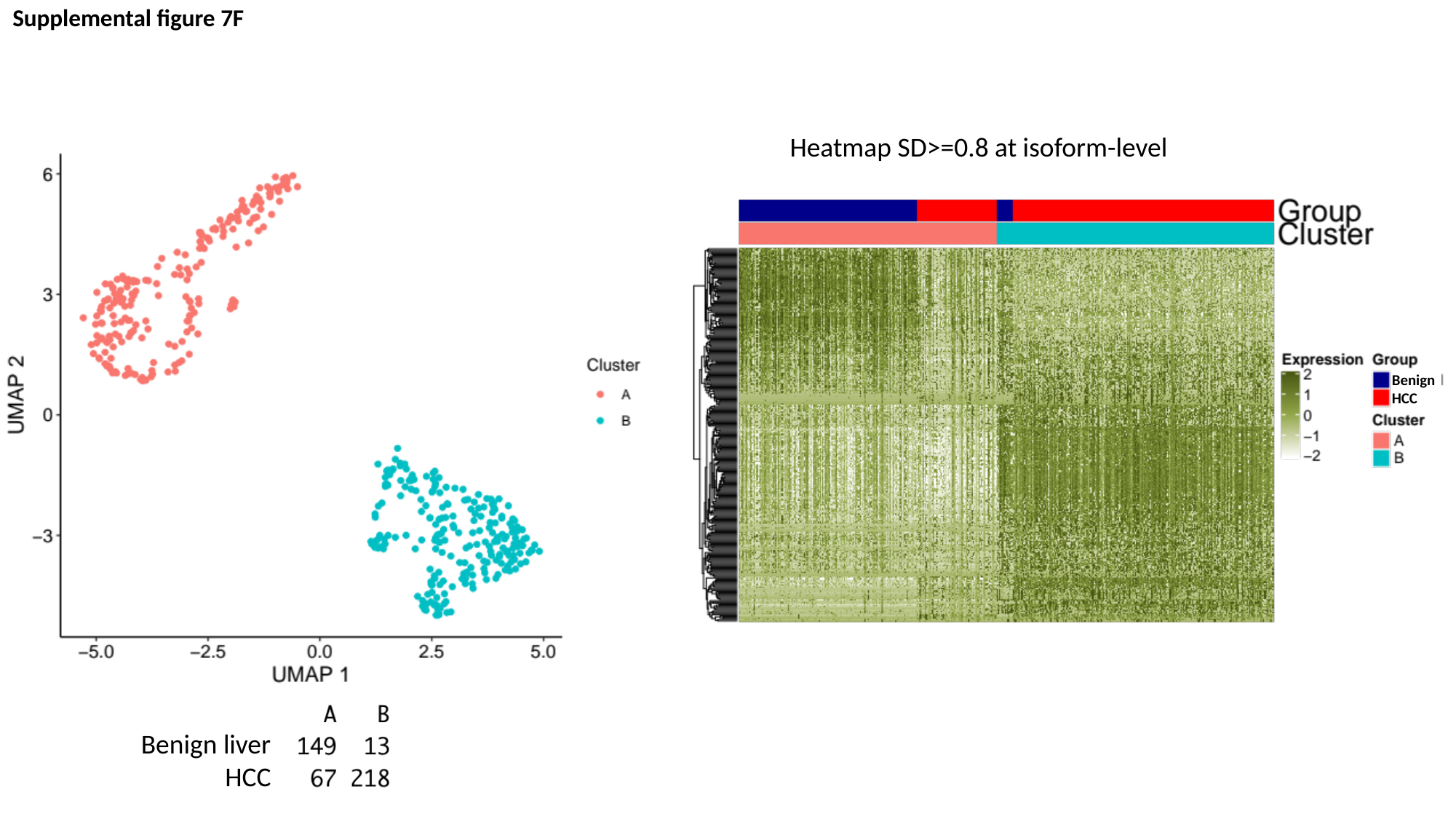

Supplemental figure 7F
Heatmap SD>=0.8 at isoform-level
Benign
HCC
Benign liver
HCC

### Slide 15
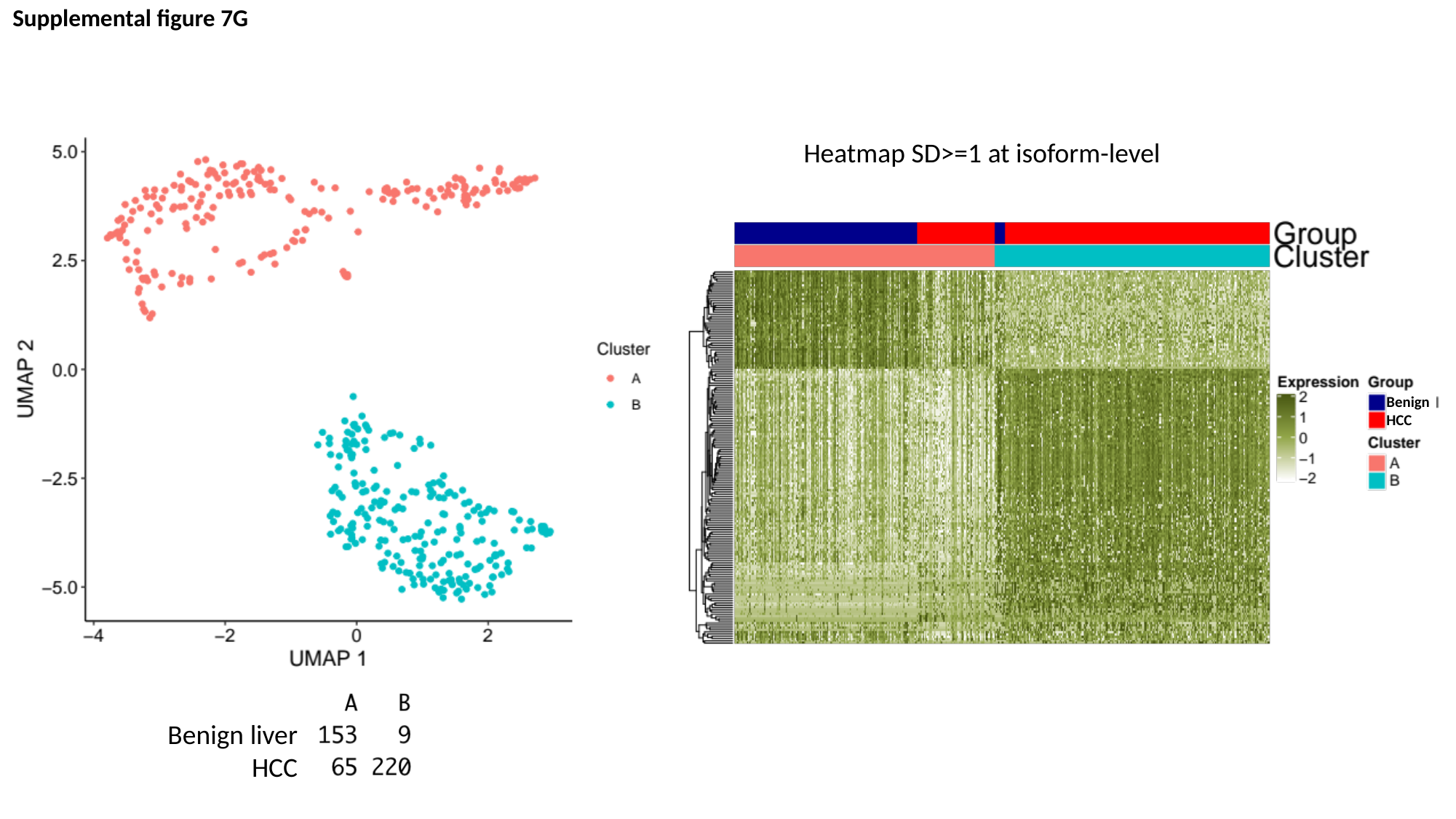

Supplemental figure 7G
Heatmap SD>=1 at isoform-level
Benign
HCC
Benign liver
HCC

### Slide 16
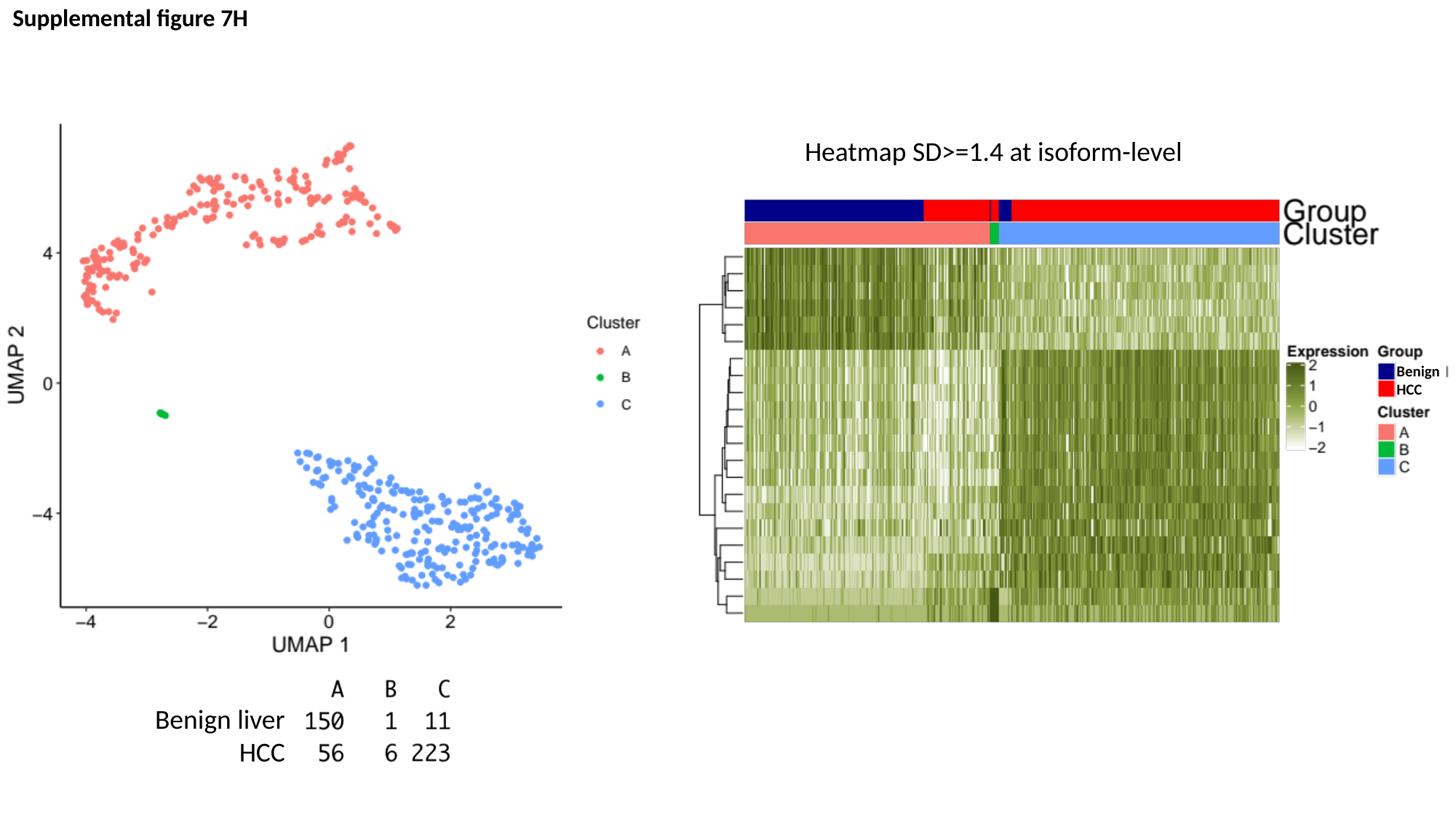

Supplemental figure 7H
Heatmap SD>=1.4 at isoform-level
Benign
HCC
Benign liver
HCC

### Slide 17
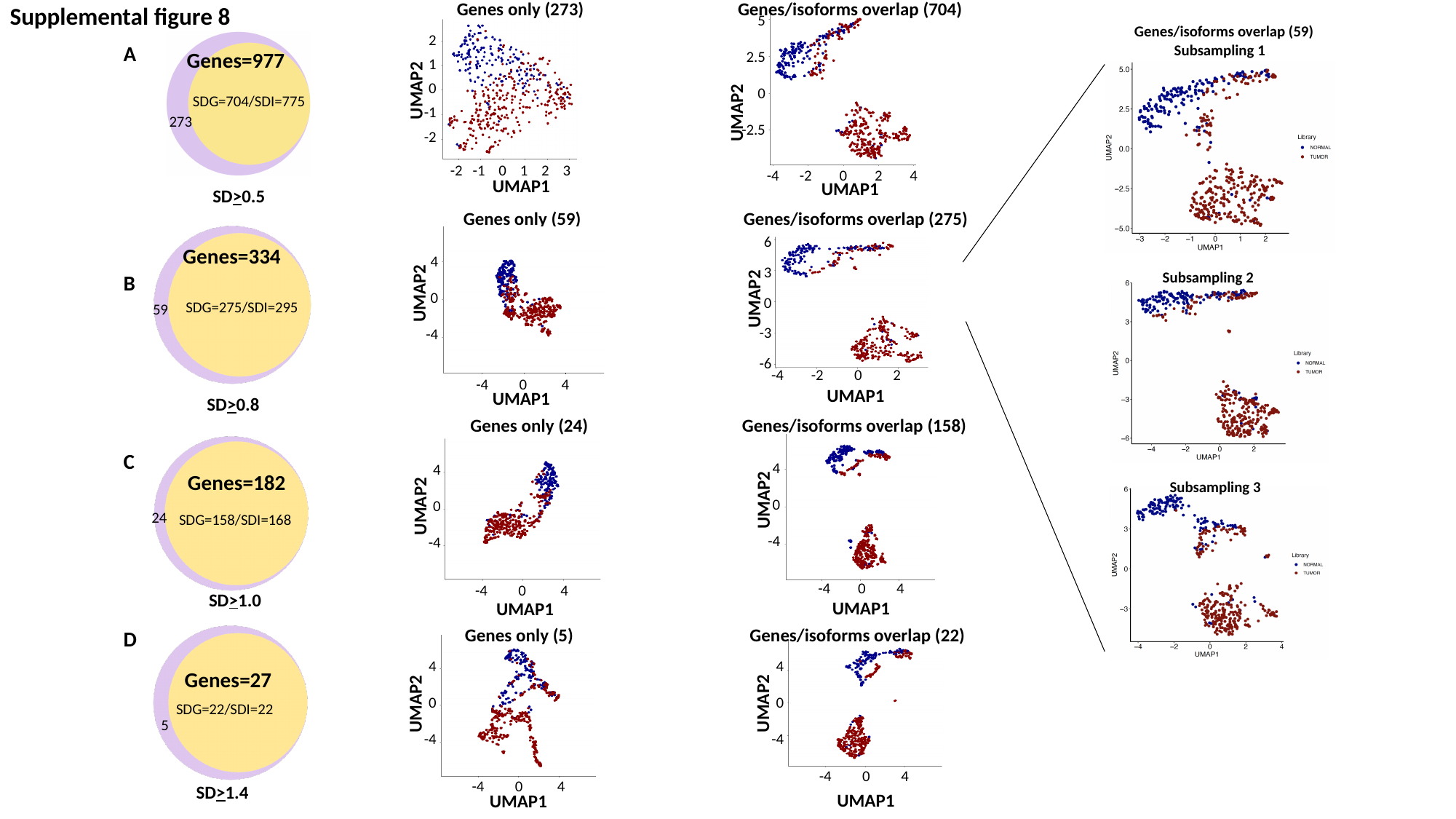

Genes only (273) Genes/isoforms overlap (704)
Supplemental figure 8
5
2.5
0
-2.5
Genes/isoforms overlap (59)
2
1
0
-1
-2
Subsampling 1
A
B
C
D
Genes=977
UMAP2
SDG=704/SDI=775
UMAP2
273
-2 -1 0 1 2 3
-4 -2 0 2 4
UMAP1
UMAP1
SD>0.5
Genes only (59) Genes/isoforms overlap (275)
6
3
0
-3
-6
Genes=334
4
0
-4
Subsampling 2
UMAP2
UMAP2
SDG=275/SDI=295
59
-4 -2 0 2
-4 0 4
UMAP1
UMAP1
SD>0.8
Genes only (24) Genes/isoforms overlap (158)
4
0
-4
4
0
-4
Genes=182
Subsampling 3
UMAP2
UMAP2
24
SDG=158/SDI=168
-4 0 4
-4 0 4
SD>1.0
UMAP1
UMAP1
Genes only (5)		 Genes/isoforms overlap (22)
4
0
-4
4
0
-4
Genes=27
UMAP2
UMAP2
SDG=22/SDI=22
5
-4 0 4
-4 0 4
SD>1.4
UMAP1
UMAP1

### Slide 18
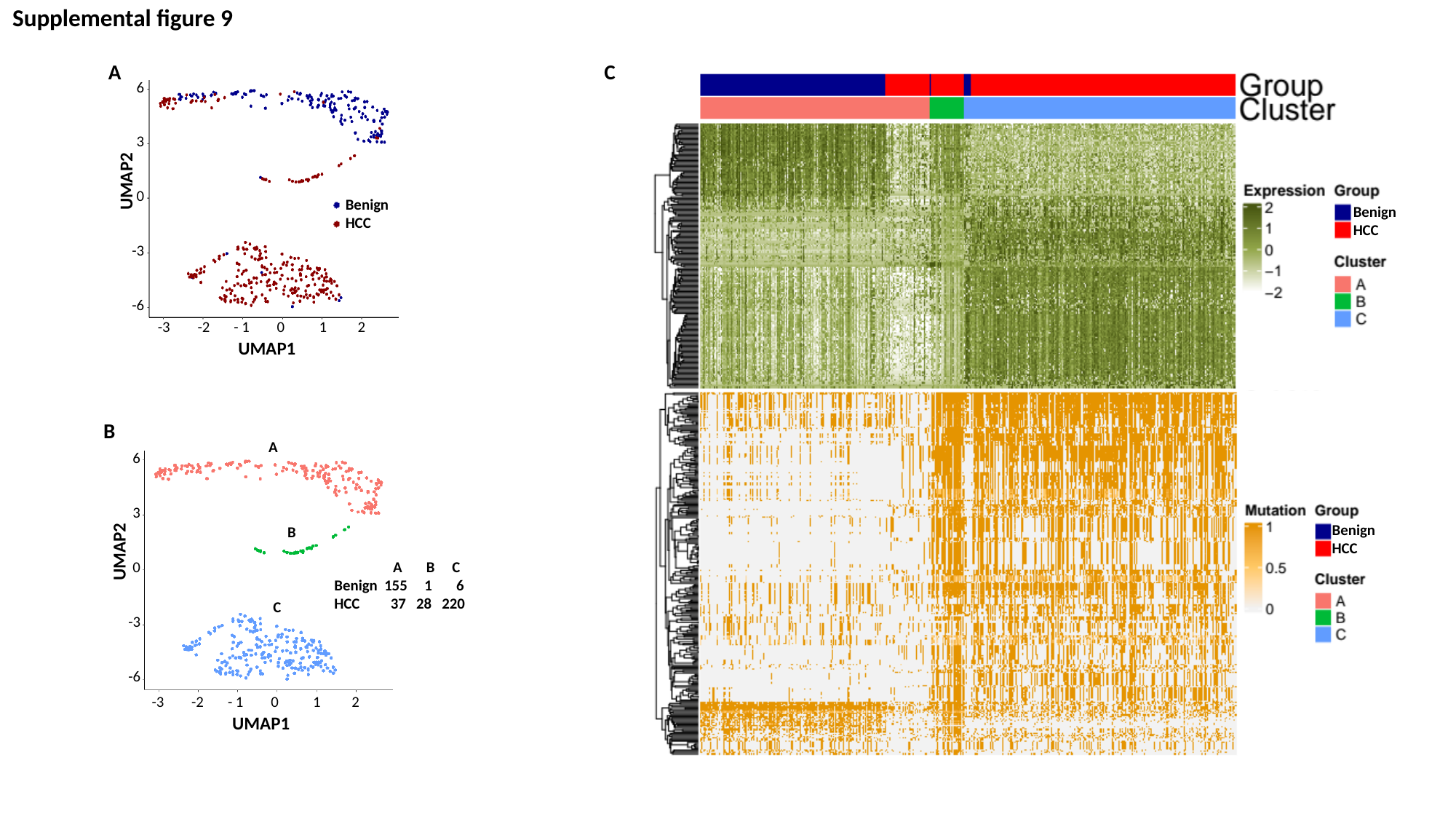

Supplemental figure 9
A
C
6
3
0
-3
-6
UMAP2
Benign
HCC
Benign
HCC
-3 -2 - 1 0 1 2
UMAP1
B
A
6
3
0
-3
-6
Benign
HCC
B
UMAP2
 A B C
Benign 155 1 6
HCC 37 28 220
C
-3 -2 - 1 0 1 2
UMAP1

### Slide 19
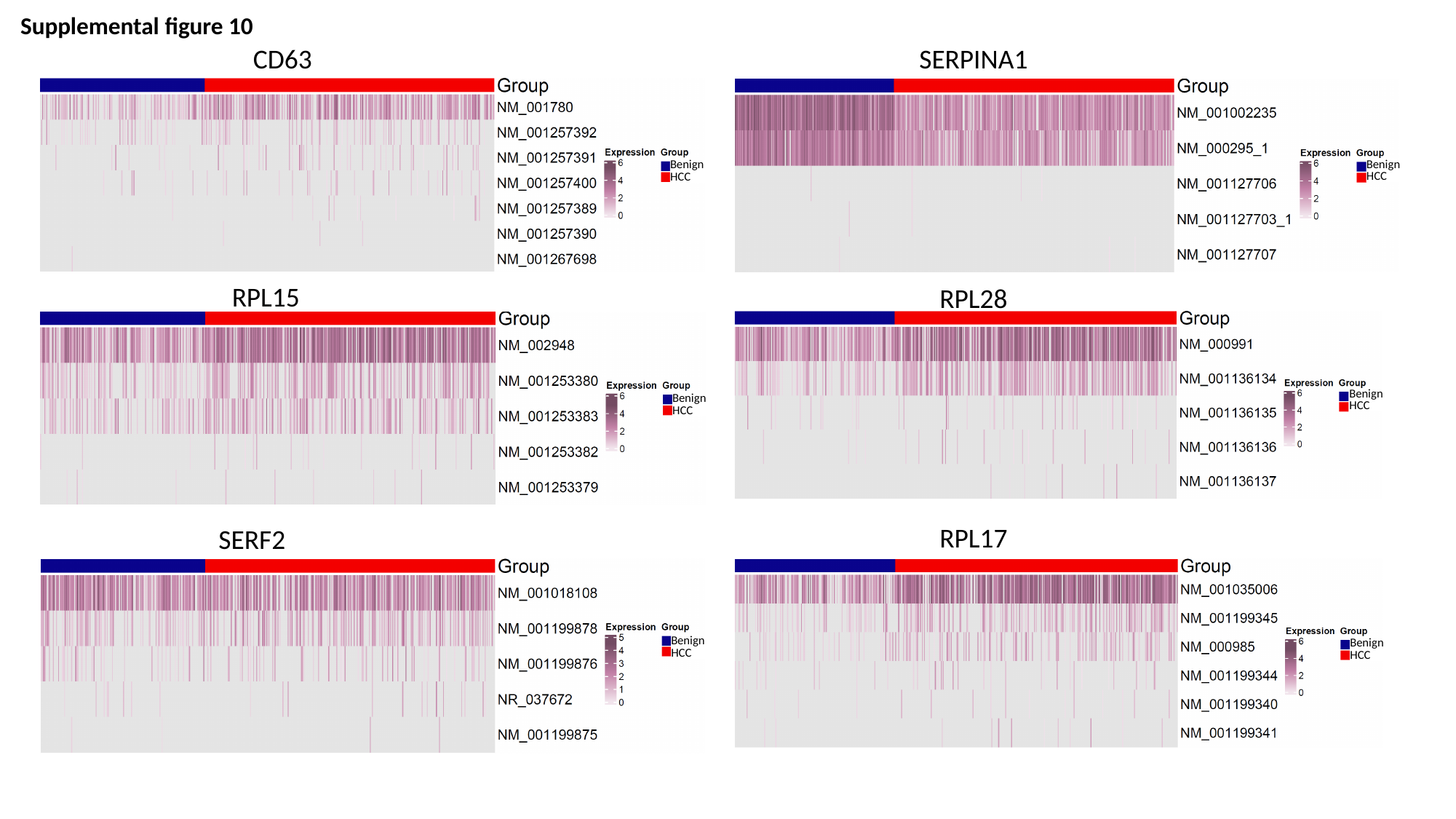

Supplemental figure 10
CD63
SERPINA1
Benign
HCC
Benign
HCC
RPL15
RPL28
Benign
HCC
Benign
HCC
RPL17
SERF2
Benign
HCC
Benign
HCC
